# Orthogonal translation initiation using the non-canonical initiator tRNA(AAC) alters protein sequence and stability *in vivo*

**DOI:** 10.1101/2021.05.25.445580

**Authors:** Andras Hutvagner, Dominic Scopelliti, Fiona Whelan, Paul R. Jaschke

**Author notes:** Correspondence to Paul R Jaschke. These authors should be regarded as joint first authors (Andras Hutvagner, Dominic Scopelliti).

## Abstract

Biological engineers seek to have better control and a more complete understanding of the process of translation initiation within cells so that they may produce proteins more efficiently, as well as to create orthogonal translation systems. Previously, initiator tRNA variants have been created that initiate translation from non-AUG start codons, but their orthogonality has never been measured and the detailed characteristics of proteins produced from them have not been well defined. In this study we created an initiator tRNA mutant with anticodon altered to AAC to be complementary to GUU start codons. We deploy this i-tRNA(AAC) into *E. coli* cells and measure translation initiation efficiency against all possible start codons. Using parallel reaction monitoring targeted mass spectrometry we identify the N-terminal amino acids of i-tRNA(AAC)-initiated reporter proteins and show these proteins have altered stability within cells. We also use structural modeling of the peptide deformylase enzyme interaction with position 1 valine peptides to interrogate a potential mechanism for accumulation of formylated-valine proteins observed by mass spectrometry. Our results demonstrate that mutant initiator tRNAs have potential to initiate translation more orthogonally than the native initiator tRNA but their interactions with cellular formyltransferases and peptide deformylases can be inefficient because of the amino acid they are charged with. Additionally, engineered initiator tRNAs may enable tuning of *in vivo* protein stability through initiation with non-methionine amino acids that alter their interaction with cellular proteases.

## Introduction

Translation initiation is an important step in expressing a functional protein and is the first crucial checkpoint in the translation process [1]. Advances in the understanding of structure and function of translational machinery, and how these biomolecules interact during the initiation process has led to an increased interest in reengineering translation initiation in order to achieve several goals within synthetic biology [2, 3]. Such goals include: enhancing the predictability and control of exogenous DNA expression, engineering biocontainment measures into genetically engineered strains, and the incorporation of non-canonical amino acids at the N-terminus of proteins [2, 4-7]. Developing orthogonal translation initiation systems may be a suitable way to achieve these goals. An ideal orthogonal translation system would maintain the most important aspects of native translation, ensuring proper functionality, but differ sufficiently from the native system to not interfere with any native expression circuits [2]. Several different orthogonal translation systems have been developed in prokaryotes, many of which focus on translation initiation as a point of engineering [3, 8-16].

A critical element of translation initiation is the interaction between the anticodon sequence of the initiator transfer RNA (i-tRNA) and the start codon, usually AUG, at the beginning of a coding sequence on an mRNA transcript. This interaction occurs within the P-site of the ribosome and is facilitated by initiation factors 1 - 3 and sequence motifs found on the mRNA (Shine-Dalgarno sequence and start codon), ribosome (regions in the 16S rRNA), and the i-tRNA (C-G motif in the anticodon stem, and the C1-A72 mismatch at the acceptor stem) [17-21]. In bacteria, the i-tRNA also carries a formylated methionine, which is incorporated exclusively at the N-terminus of the growing polypeptide chain [22]. Methionine residues lacking formylation are also incorporated throughout the growing polypeptide chain, however these amino acids, along with any other amino acids found internally within a protein are delivered to the growing peptide chain by elongator tRNAs [22].

All transfer RNAs share similar structural elements, including the D-loop, anticodon loop, the TΨC-loop and acceptor stem (Fig. 1). These structures stabilize the tRNA during translation and house nucleotide motifs which are vital during tRNA aminoacylation and P-site binding. For instance, it has been shown that the unpaired nucleotide in position 73 on the tRNA, along with nucleotide pairs at 1:72, are vital synthetase recognition elements in most tRNA species (Fig. 1). Furthermore, the variable loop, the D-loop, and the anticodon also contain important aminoacyl-synthetase recognition elements, which act to ensure proper aminoacylation of the tRNA [11, 23-26].

**Figure 1.**
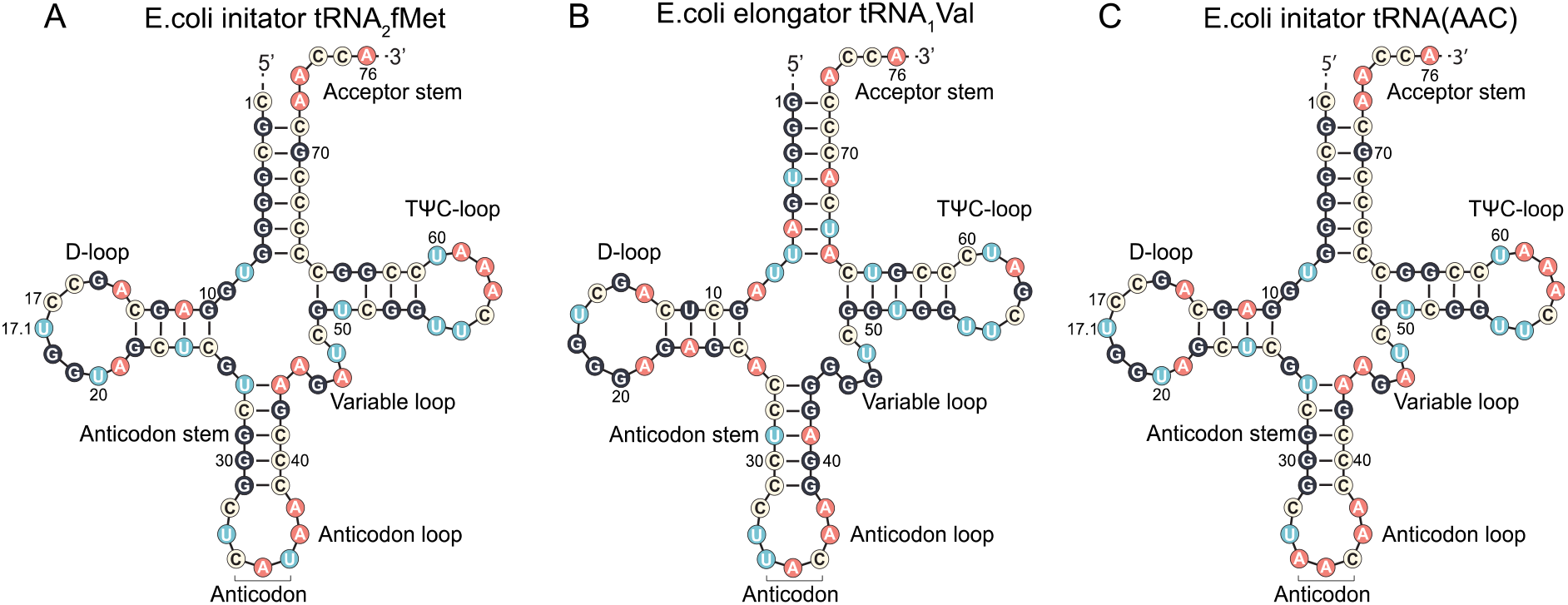
Similarities and differences between *E. coli* initiator and elongator tRNAs. *Escherichia coli* initiator tRNA_2_^fMet^ (left), the *E. coli* elongator tRNA_1_^Val^ (middle) depicted, and the mutant *E. coli* initiator tRNA(AAC) (right).

Although i-tRNAs and elongator tRNAs share an abundance of structural similarities, several sequence motifs unique to the i-tRNA enable this molecule to perform a unique role in translation initiation. Firstly, i-tRNAs possess a mismatch between nucleotides C1 and A72 which is a recognition element for methionyl-tRNA transformylase (FMT) [27]. After methionine is charged to i-tRNA in bacteria, the FMT enzyme formylates the bound methionine [27, 28]. This fMet residue is essential for subsequent interaction between i-tRNA^fMet^ and initiation factor 2 prior to recruitment to the P-site of the 30S complex, and some evidence suggests initiation cannot occur without a formylated amino acid bound to i-tRNA [29]. Secondly, a highly conserved 3G-C motif in the anticodon stem plays a vital role in P-site binding through interactions with the 16S rRNA of the ribosome and initiation factor 3 [21, 30].

Recently, it has been observed that to some degree in nature, translation can initiate from the majority of non-canonical start codons, although at significantly lower rates than the canonical AUG, GUG, and UUG start codons [31, 32]. This sets a strong precedent for the use of mutant i-tRNAs which couple with non-canonical start codons, as it seems that simply a strong anticodon-codon interaction may be enough to pass the first checkpoint of translation initiation in prokaryotes. Previous studies have used engineered non-canonical i-tRNAs to initiate translation from non-canonical start codons [3, 8, 10, 11, 15, 16], however, they did not characterise how orthogonal these initiators might be to other potential start codons.

In this study, we created and characterized a mutant i-tRNA with AAC anticodon in place of the canonical CAU anticodon. We measured the efficiency of translation initiation of this i-tRNA(AAC) against all the 63 possible non-AUG start codons and found it is more orthogonal than the native i-tRNA(CAU). Using targeted mass spectrometry methods we identify valine and formylated-valine at the N-terminus of i-tRNA(AAC) initiated proteins, and measure their stability *in vivo*. Finally, we computationally model the interaction between peptide deformylase and formyl-valine peptides to propose a mechanism explaining potential deficiencies in the removal of the formyl group from i-tRNA(AAC)-initiated proteins.

## Results

### Measuring translation initiation efficiency of i-tRNA(AAC) against all possible start codons

To create a new mutant initiator tRNA within *Escherichia coli* we first searched for an anticodon with low or no utilization within *E. coli*. Using the Genomic tRNA Database, we found that no elongator tRNA carrying a cognate AAC anticodon is present in *E. coli* [33]. To determine if it would be possible to express a functional i-tRNA with this anticodon, we built a plasmid containing a modified *metY* gene to express a mutant version of the canonical i-tRNA_2_^fMet^. The anticodon region of *metY* from MG1655 strain was changed from CAT to AAC and inserted into pULTRA plasmid to make pULTRA::*tac*-*metY*(AAC) (Fig. 2A). A control version of this plasmid (pULTRA::*tac*-Empty) lacking *metY*(AAC) gene was also made (Fig. 2B).

**Figure 2.**
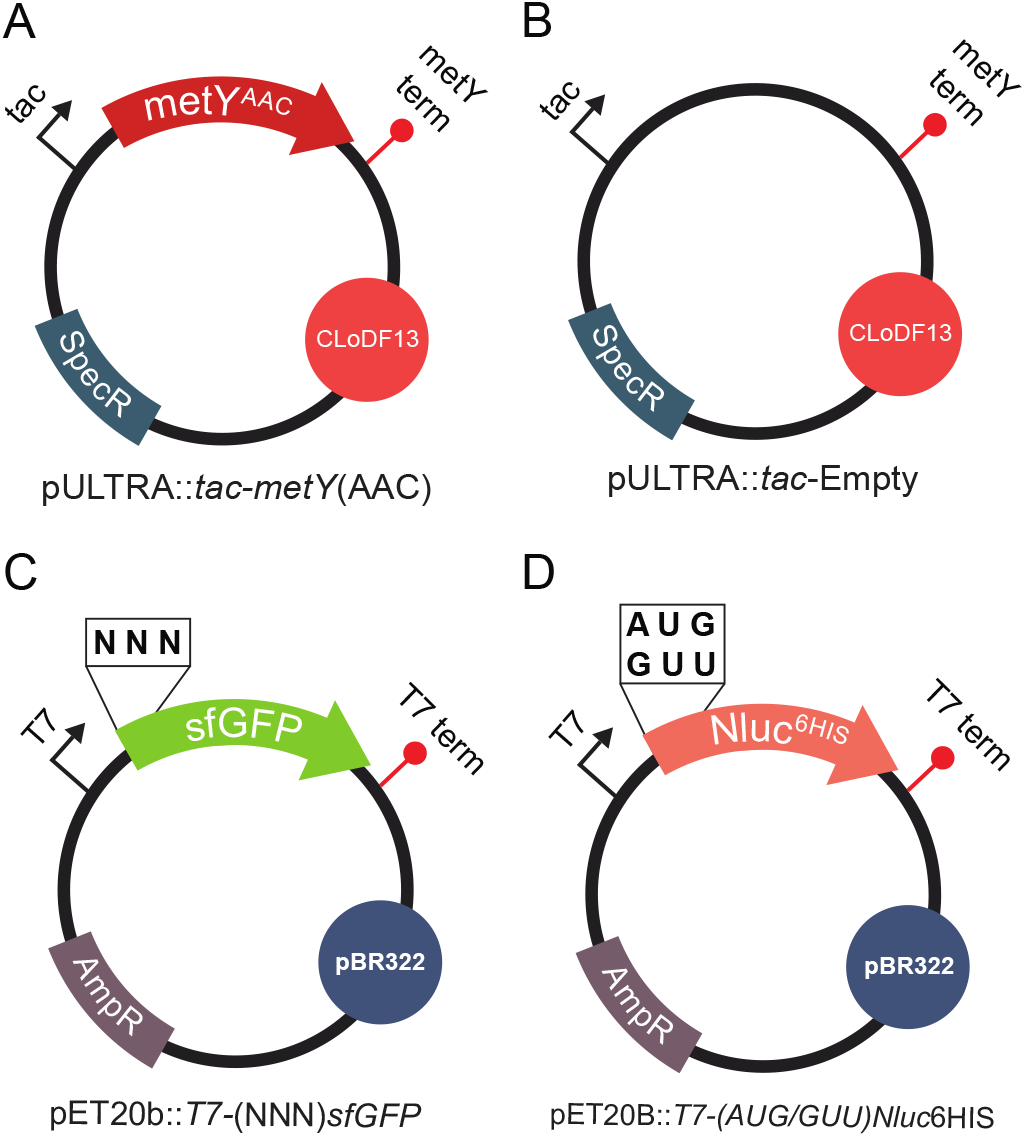
Plasmids used in this study. (A) pULTRA::*tac*-*metY*(AAC) expresses the i-tRNA(AAC) anticodon mutant and contains a medium copy ClodF13 origin of replication, spectinomycin resistance gene, inducible *tac* promoter, and *metY* terminator. (B) pULTRA::*tac*-Empty control plasmid is identical to pULTRA::*tac*-*metY*(AAC) but lacks *metY* gene. (C) pET20b:*T7-*(NNN)*sfGFP* expresses the sfGFP reporter with altered start codons and contains a high copy pBR322 origin of replication, ampicillin resistance gene, T7 promoter, and T7 terminator. (D) pET20b:*T7-*(AUG/GUU)*Nluc*6His expresses the luciferase Nluc with altered start codons, C-terminal 6xHis tag and contains a high copy pBR322 origin of replication, ampicillin resistance gene, T7 promoter, and T7 terminator. All plasmids were designed to have compatible origins of replication and selectable markers.

In order to measure the translation initiation efficiency of mutant i-tRNA(AAC), we used a previously described superfolder GFP (sfGFP) fluorescent reporter library to determine how efficiently i-tRNA(AAC) initiates translation from all 64 start codons [31]. We transformed *E. coli* strain BL21(DE3)pLysS with: (1) the mutant i-tRNA plasmid harboring the modified tRNA *metY* gene with an AAC anticodon region (pULTRA::*tac*-*metY*(AAC) (Fig. 2A) or pULTRA::*tac*-Empty (Fig. 2B), and (2) a set of reporter plasmids harboring sfGFP with all possible start codons (pET20b::T7-(NNN)sfGFP) (Fig. 2C).

To investigate how the presence of i-tRNA(AAC) within the *E. coli* cell affects the translation efficiency of (NNN)sfGFP, we analyzed the bulk cell and population fluorescence (Arb.U.) of strains harboring the pULTRA::*tac*-*metY*(AAC) and pET20b::T7-(NNN)sfGFP plasmids, under inducing conditions and compared to cells containing pULTRA::*tac*-Empty. Relative fluorescence of cells containing the pET20b::T7-(NNN)sfGFP reporters and the control plasmid pULTRA::*tac*-Empty were broadly similar to those reported in Hecht, Glasgow [31], with the strongest initiation events occurring from canonical start codons AUG, GUG, and UUG (Fig. 3). In contrast, when cells contained pULTRA::*tac*-*metY*(AAC), we observed an increase in translation initiation from GUU start codon of 8-fold over that seen in cells containing pULTRA::*tac*-Empty (Fig. 3). Similarly, measurements of the same cultures using flow cytometry showed unimodal cell populations for all start codons and a 10-fold increase in fluorescence from GUU start codon (Fig. S1).

**Figure 3.**
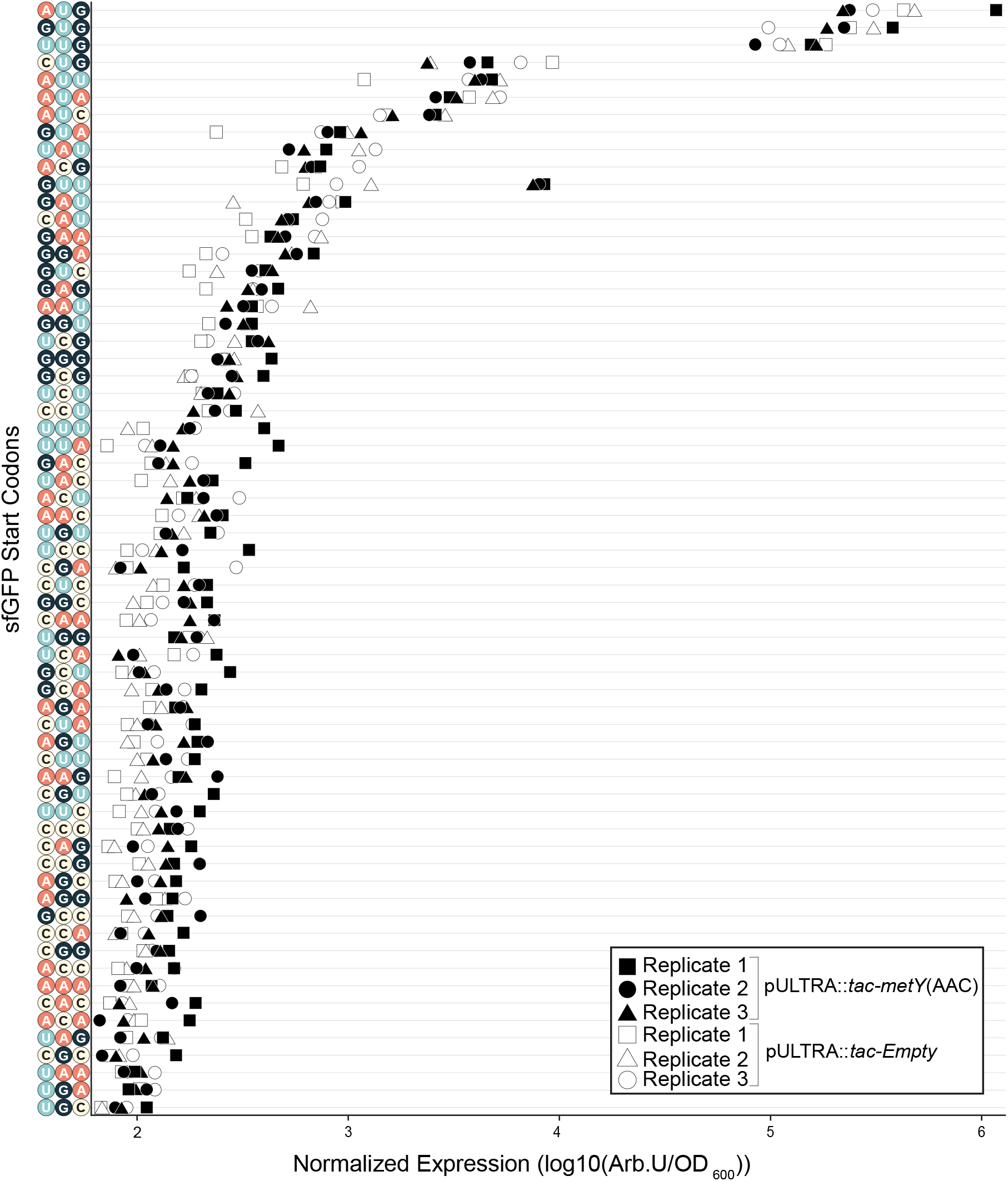
Efficiency of translation initiation of all sfGFP variants using the mutant i-tRNA(AAC). Normalized bulk cell fluorescence (Arb. U./OD_600_) of all sfGFP start codon variants was captured from cells grown to mid-log phase (0.6 OD_600_) in the presence of the i-tRNA(AAC) induced with 1 mM IPTG for 1-hour prior to measurement.

To determine if there were any other significantly differentially initiating start codons as a result of i-tRNA(AAC), we repurposed the differential expression analysis for sequence count data 2 (DESeq2) method normally used for transcriptomics [34]. Considering that the fluorescence data gathered in this study was in the form of integer count data and the number of replicates allowed the statistical models to hold true, we reasoned that DESeq2 was appropriate for this analysis. By designating the set of strains containing the pULTRA::*tac*-Empty and pULTRA::*tac*-*metY*(AAC) plasmids as sample conditions, we analyzed the efficiency of i-tRNA(AAC) initiation against all 64 possible start codons. DESeq2 analysis of the bulk fluorescence and flow cytometry data showed only start codon GUU was significantly (p-value <0.01) differentially expressed in the presence of i-tRNA(AAC) (Fig. 4). This result shows that the translation initiation enhancing effect of i-tRNA(AAC) is limited to its cognate start codon GUU.

**Figure 4.**
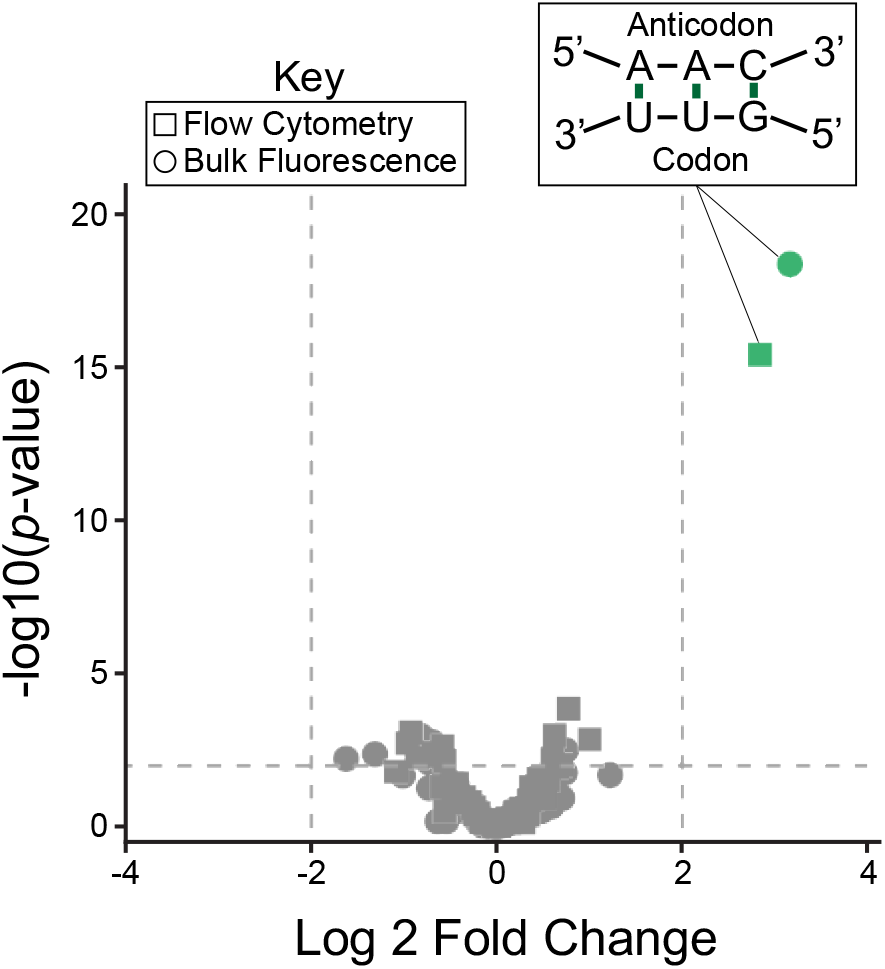
DESeq2 analysis of translation initiation from i-tRNA(AAC) shows only translation initiation from cognate GUU start codon increased by mutant i-tRNA(AAC). DEseq2 analysis was used to compare the normalized bulk cell fluorescence and flow cytometry fluorescence (Arb. U./OD_600_) of cells containing i-tRNA(AAC) and sfGFP reporter library, to control cells containing pULTRA:*tac*-Empty and sfGFP reporter library.

With the orthogonality of i-tRNA(AAC) within *E. coli* established, we next wanted to determine whether it had any effects on cellular fitness. We cultured BL21(DE3)::pLysS cells carrying either pULTRA::*tac*-*metY*(AAC) or pULTRA::*tac*-Empty plasmids under either inducing or non-inducing conditions, and measured both growth rate and final OD_600_ for both. The results of this analysis showed that there was a significant decrease in the growth rate of i-tRNA(AAC) expressing cells (p<0.05), while no significant changes to maximum cell density were seen (Fig. 5). The uninduced condition showed no significant changes to maximum cell density or growth rate.

**Figure 5.**
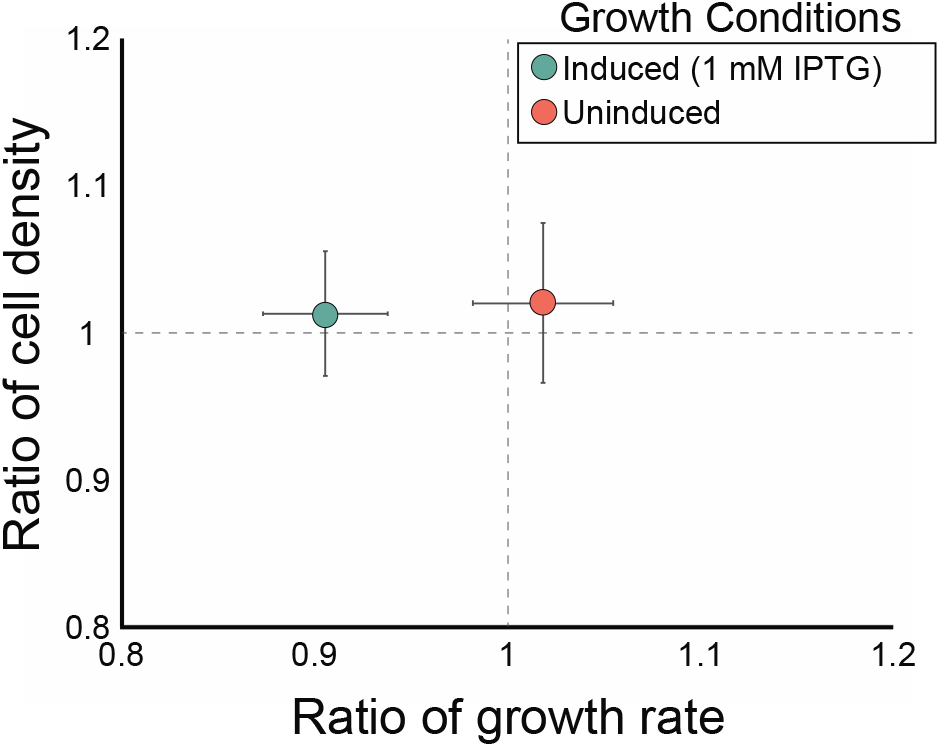
Fitness analysis of a strain harboring pULTRA::*tac*-*metY*(AAC). Ratios of growth rate and maximum cell density were determined for strains harboring pULTRA::*tac*-*metY*(AAC) and compared to pULTRA::*tac*-Empty control. All strains were grown in LB in the presence of spectinomycin and chloramphenicol in biological triplicate. 1 mM IPTG was added to the induced condition, whereas equivalent amounts of sterile water was added to the uninduced condition. Error bars represent one standard deviation.

### Identification of the amino acid incorporated onto the N-terminus of proteins expressed using i-tRNA(AAC)

Since there is no initiator tRNA within *E. coli* that carries the AAC anticodon, we next wanted to confirm which amino acid was aminoacylated to i-tRNA(AAC). To identify the N-terminal amino acid used to initiate translation from GUU start codons, we created two new C-terminally His-tagged NanoLuc luciferase (Nluc) reporter plasmids containing GUU and AUG start codons: pET20b:T7-(GUU)*Nluc*6HIS and pET20b:T7-(AUG)*Nluc*6HIS (Fig. 2D). To express the (GUU)Nluc protein while also expressing i-tRNA(AAC), along with appropriate controls, we grew BL21(DE3)pLysS containing pET20b:T7-(GUU)*Nluc*6HIS or pET20b:T7-(AUG)*Nluc*6HIS along with either pULTRA::*tac*-*metY*(AAC) or pULTRA::*tac*-Empty. Next, we Ni-NTA purified the Nluc proteins, digested with trypsin, and subjected the resulting peptides to mass spectrometry.

The mass spectrometry approach parallel reaction monitoring (PRM) was used to identify the amino acid incorporated onto the N-terminus of the Nluc reporters. The main benefit of using PRM is that this method allows us to predefine specific targets, providing a high degree of selectivity and sensitivity by excluding undesirable signals. The isolation list used to specify the targeted peptides in this PRM analysis included several internal Nluc peptides and the 20 possible amino acids with and without formylation at the N-terminus (File S1). As expected, in cells carrying pET20b:T7-(AUG)*Nluc*6HIS and pULTRA::*tac*-Empty plasmids, we found that N-terminal peptides of Nluc proteins carried methionine exclusively (807.8854 *m/z*) (Fig. 6A). Similarly, in cells carrying pET20b:T7-(GUU)*Nluc*6HIS and pULTRA::*tac*-Empty plasmids, all N-terminal peptides had only methionine (Fig. 6B), supporting previous results on initiation from non-canonical start codons [31].

**Figure 6.**
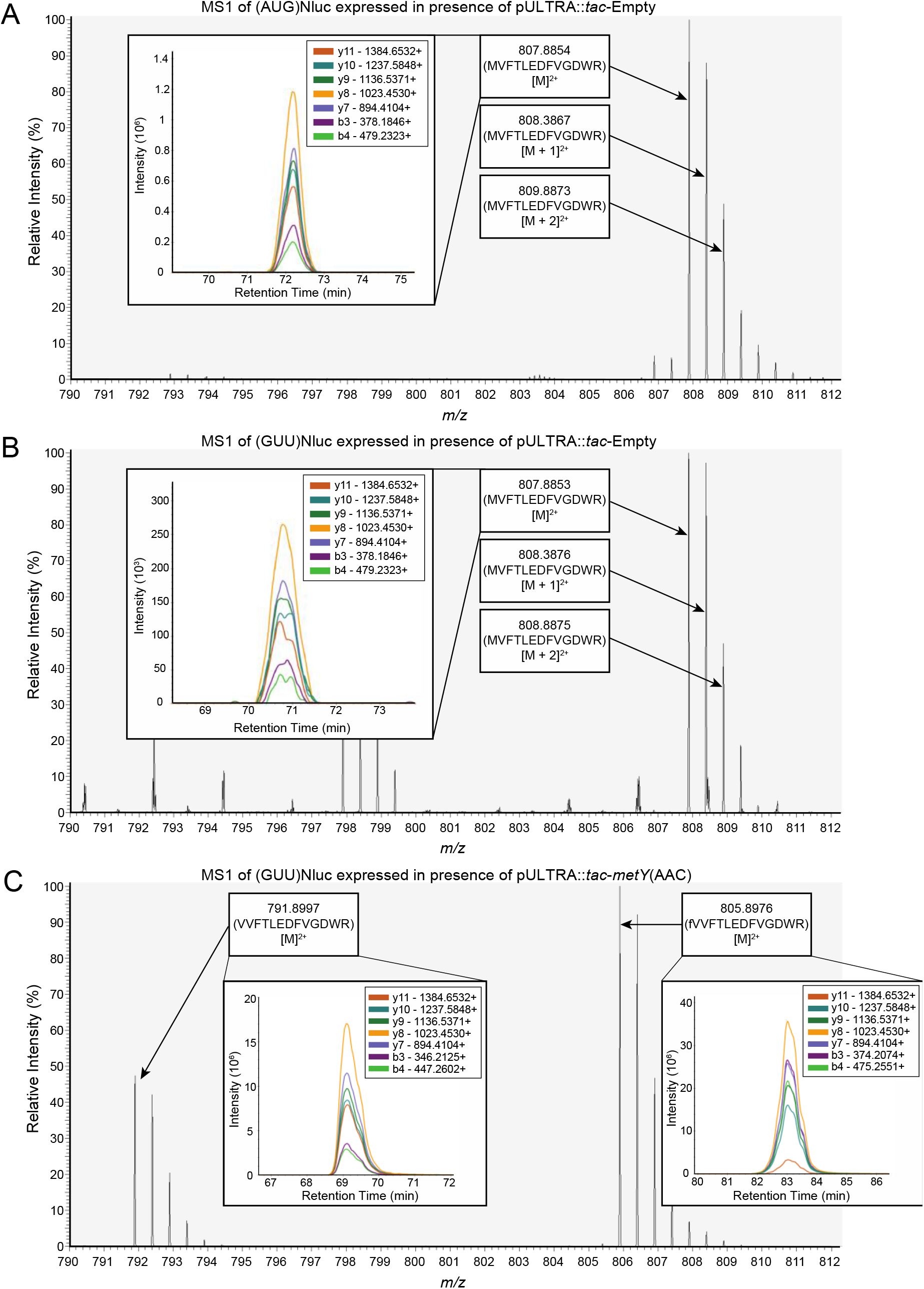
N-terminal amino acid residue identification of Nluc using targeted mass spectrometry. Nluc-6His reporter proteins carrying either AUG or GUU start codons were Ni-NTA purified from three different BL21(DE3)pLysS strains possessing either the pULTRA::*tac*-*metY*(AAC) or pULTRA::*tac*-Empty. (A) MS1 scan during retention time range of 68-73 minutes of Nluc reporter protein expressed in BL21(DE3)pLysS (pULTRA::*tac*-Empty; pET20b:T7-(AUG)*Nluc*6HIS). Labelled precursor ion peaks (807.8854 *m/z*) consistent with an N-terminal peptide bearing methionine (MVFTLEDFVGDWR) and their respective isotopes at the N-terminus are shown. Inset shows MS2 y^-^ and b^-^ series product ion signals used to confirm amino acid sequence of N-terminal peptide. (B) MS1 scan of Nluc during retention time range of 68-73 minutes of Nluc reporter protein expressed in BL21(DE3)pLysS (pULTRA::*tac*-Empty; pET20b:T7-(GUU)*Nluc*6HIS). Labelled precursor ion peaks (807.8853 *m/z*) consistent with an N-terminal peptide bearing methionine (MVFTLEDFVGDWR) and their respective isotopes at the N-terminus are shown, with no evidence of a peptide with an N-terminal valine residue present. Inset shows MS2 y^-^ and b^-^ series product ion signals used to confirm amino acid sequence of N-terminal peptide. (C) MS1 scan between retention times 68-84 minutes of Nluc reporter protein expressed in BL21(DE3)pLysS (pULTRA::*tac*-*metY*(AAC); pET20b:T7-(GUU)*Nluc*6HIS). Labelled precursor ion peaks (791.8997 *m/z* and 805.8976 *m/z*) are consistent with N-terminal peptides bearing either valine (VVFTLEDFVGDWR) or formylated valine (fVVFTLEDFVGDWR). Insets show MS2 y^-^ and b^-^ series product ion signals used to confirm amino acid sequence of N-terminal peptide.

In contrast, cells carrying both pET20b:T7-(GUU)*Nluc*6HIS and pULTRA::*tac*-*metY*(AAC) plasmids showed two different precursor ions (791.8997 *m/z* and 805.8976 *m/z*) that were absent in the (AUG)Nluc samples (Fig. 6C). These two new precursor *m/z* values are consistent with N-terminal peptides bearing a valine and a formylated valine residue on the N-terminus of the (GUU)Nluc protein and was confirmed by subsequent MS2 analysis (Fig. 6C, insets). Interestingly, in addition to valine bearing N-terminal peptides, methionine (but not formylated-methionine) bearing N-terminal peptides were also identified, suggesting a heterogenous population (Fig. S2).

### Determination of the stability of (GUU)Nluc proteins within *E. coli*

With the knowledge that the majority of the N-terminal residues of (GUU)Nluc were either valine or formylated valine, we wanted to measure whether there were any effects on the stability of the reporter protein from this altered N-terminal amino acid. To measure *in vivo* protein stability of the Nluc reporters using the enzymatic activity of the protein (luminescence) as a proxy for protein levels, we spun down and washed overnight cultures of BL21(DE3)pLysS strains containing combinations of the reporter and i-tRNA expressing plasmids, pulsed them with IPTG for 1 hour, followed by growth in repressing conditions for 12 hours. Early time points in the luminescence dataset showed considerably less luminescence from reporter proteins expressed from a GUU start codon, as opposed to reporters expressed from the native AUG start codon. These findings are in agreement with the findings of the bulk fluorescence and flow cytometry, which showed that translation initiation of sfGFP from GUU start codons in the presence of the mutant i-tRNA(AAC) is several orders of magnitude less efficient than translation initiation from AUG codons by native i-tRNA-fMet (Fig. 3).

We observed a relatively steady decrease in luminescence over the 12 hour time course for (GUU)Nluc expressed in the presence of i-tRNA(AAC) and the native (AUG)Nluc expressed in the absence of i-tRNA(AAC) (Fig. 7A). We observed very low signal and no change in measurable levels of luminescence from the (GUU)Nluc reporter expressed in the absence of i-tRNA(AAC), over the 9 hour time-course (Fig. 7A).

**Figure 7.**
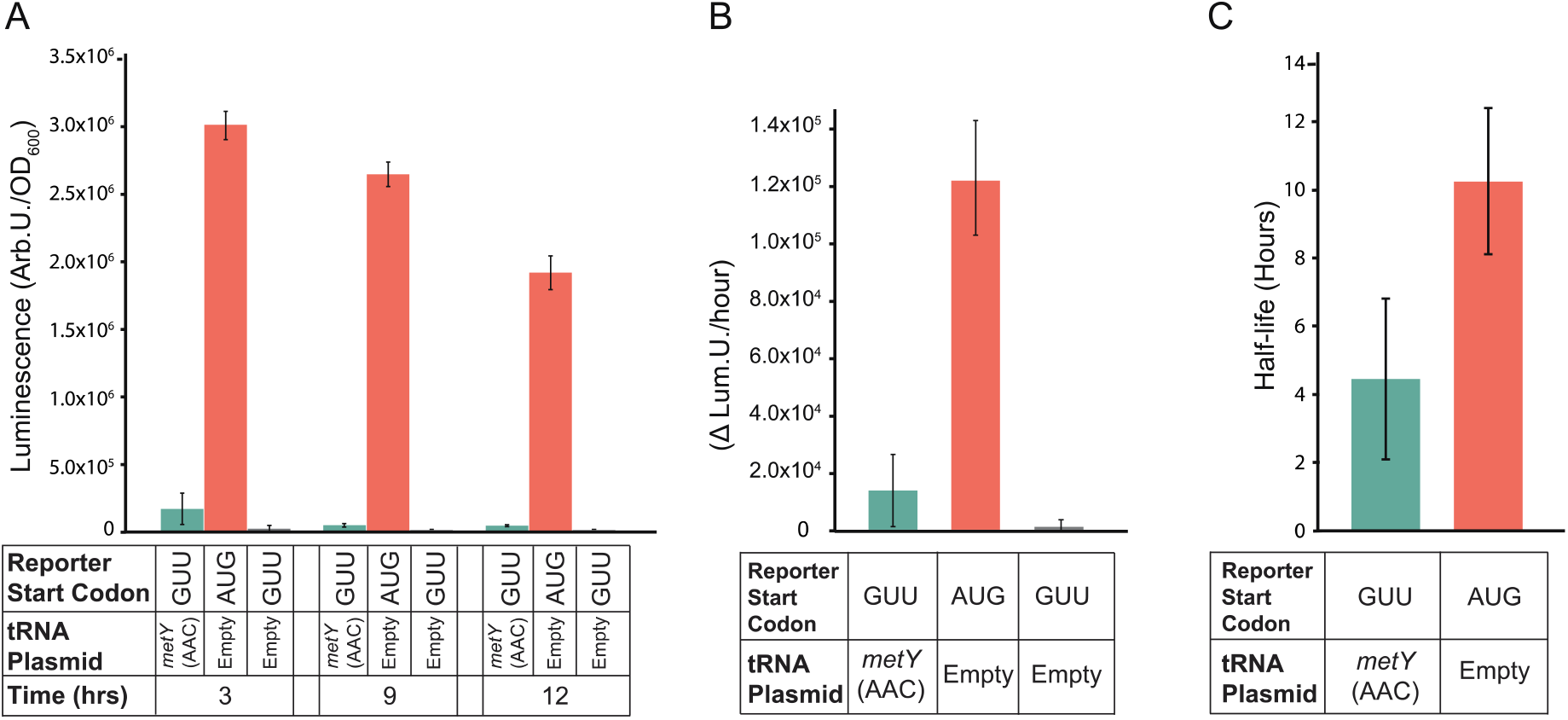
*In vivo* stability of Nluc proteins expressed in the presence and absence of i-tRNA(AAC). Nluc luminescence was analysed from BL21(DE3)pLysS strains grown in biological triplicates. Saturated cultures of each strain were back diluted to 0.1 OD_600_ and grown for 2 hours prior to a 1-hour pulse induction with 1 mM IPTG. Cells were then washed and resuspended in repressing conditions (LB, 2% glucose) and grown for 12 hours, with luminescence measured every 3 hours with the H1 Hybrid Multi-Mode Reader at 100 gain setting. (A) Normalized luminescence from *E. coli* BL21(DE3)pLysS cells harbouring either pULTRA::*tac*-*metY*(AAC)/ pET20b:T7-(GUU)*Nluc*6HIS (green), pULTRA::*tac*-Empty/ pET20b:T7-(AUG)*Nluc*6HIS (orange), or pULTRA::*tac*-Empty/ pET20b:T7-(GUU)*Nluc*6HIS plasmids (grey) at 3, 9, and 12 hours post induction. (B) Calculated rates of luminescence decrease per hour from 3- and 12-hours luminescence readings post induction. (C) Half-lives of Nluc calculated using normalized luminescence from 3- and 12-hours post induction.

The rate of degradation was found to be approximately 8 times faster in the native (AUG)Nluc samples compared to the (GUU)Nluc samples (Fig. 7B), but the half-life of (GUU)Nluc was only 4.5 hours compared to 10.3 hours for (AUG)Nluc (Fig. 7C) indicating proteins expressed using i-tRNA(AAC) were significantly less stable than from the (AUG)Nluc control.

### Comparing formyl-valine and formyl-methionine tripeptides as substrates for peptide deformylase

Given that we detected a strong signal of formylated valine peptides from (GUU)Nluc expressed in the presence of i-tRNA(AAC) (Fig. 6C), but no formylated methionine signals from (GUU)Nluc (Fig. 6B), and only weak signals in (AUG)Nluc in the absence of i-tRNA(AAC) (Fig. S3), this suggested to us that the interaction between formylated valine and the enzyme responsible for removing the formyl group was altered from that of formyl methionine. We next wanted to understand the reason for this apparent difference. In *E. coli* the formyl group is normally removed from the N-terminal methionine by the essential enzyme peptide deformylase. Thus, we investigated the relative binding stability of XAM peptides (where X is any residue except proline) to the *E. coli* enzyme peptide deformylase *in silico* using a molecular mechanics approach. Here, we have studied the Ni^2+^ bound enzyme structure (pdb code: 1bs6) as this is has been shown to be an active deformylase [35] and has been defined bound to partial substrate peptide MAS [36]. Molsoft ICM-Pro software was used to calculate the relative difference in binding free energy (ΔΔG) between mutant and wildtype complexes, computed from the sum of van der Waals attraction and repulsion, electrostatic interactions, change in entropy, hydrogen bond formation and solvent interactions using the Internal Coordinate Force Field (ICFF) [37]. A Monte Carlo simulation of side chain repacking in the vicinity of the mutated residue was performed to optimise the lowest energy mutant conformation and binding interaction while maintaining the geometry of the peptide backbone [38-40].

A negative ΔΔG indicates a higher stability interaction; a positive ΔΔG indicates a lower stability interaction, while a ΔΔG of ≥2.5 kcal/mol is considered a significant difference [38]. Thus, peptides with Q, V, S, N, E, C, W, A, G or D at position 1 (pdb code 1bs6, chain D [36] are predicted from this analysis to have significantly lower stability binding in complex with *E. coli* peptide deformylase (1bs6, chain A) (Table 1). In support of this model, it has previously been demonstrated that this enzyme cannot hydrolyse a formylated tetrapeptide with Ala at position 1 (fAGSE) [41].

**Table 1.** Relative protein:protein binding free energy ΔΔG for mutant peptides XAS.

| Amino Acid | $\Delta\Delta G$ |
| --- | --- |
| D | 6.06 |
| G | 5.75 |
| A | 4.51 |
| W | 4.07 |
| C | 3.73 |
| E | 3.41 |
| N | 3.37 |
| S | 3.21 |
| V | 2.89 |
| Q | 2.75 |
| R | 2.25 |
| H | 2.05 |
| Y | 2.01 |
| T | 1.94 |
| F | 1.92 |
| I | 1.90 |
| L | 1.75 |
| K | 1.31 |
| M | 0.00 |

$\Delta\Delta G$ (kcal/mol) is the difference in Gibbs free energy ( $\Delta G$ ) of mutant peptide binding relative to wild type peptide, $\Delta\Delta G = \Delta G^{\text{mutant}} - \Delta G^{\text{wt}}$ .

A more detailed prediction of possible binding poses was performed by docking models of formylated substrates fMAS, fVAS and fAAS into the Ni-bound active site of peptide deformylase (1bs6, chain A). The docking score (Edoc) represents a composite score where the lower the score, the better the binding, and is calculated from a combination of hydrogen bond grid energy (Egb), electrostatic grid potential (Ege), hydrophobic potential (Egs) and van der Waals interaction (Egv) (Tables S1-S3). The best scoring example of each peptide indicates a binding preference in the order fMAS>fVAS>fAAS (Edoc scores) (Table 2). This correlated with Egv and Egs scores, which demonstrated the highest variance between mutant peptides for the best poses, indicating that van der Waals and hydrophobic interactions are likely the basis of substrate selectivity. Docking was performed in triplicate; and the most negative scoring docked peptides show very limited variance in backbone architecture proximal to the active site (Fig. 8A), with the fMAS peptide having the formyl carbonyl carbon coordinated in proximity to the catalytic nucleophile W1 (3.0 Å), the backbone amide hydrogen of L91 (2.3 Å), and the active site Ni atom (3.4 Å) (Fig. 8B), consistent with the mechanism of peptide deformylase formyl hydrolysis proposed by Becker et al. [36] and modelled by Wu et al. [42]. The highest scoring docked fVAS peptide shows a similar orientation, with the formyl carbonyl carbon more closely associated with W1 (2.9 Å) and Ni (3.0 Å), and slightly further from L91 (2.7 Å) (Fig. 8C). Thus, no profound differences in the active site localisation of the formyl group are evident, likely indicating that this binding event would be competent for catalysis. The molecular determinants of specificity are clearer when rendered as a 2-dimensional interaction diagram. A decrease in hydrophobic interactions is evident for fVAS, becoming more pronounced for fAAS relative to fMAS (Fig. 9). The progressive decrease in accessible surface area for the side chain at position 1 may explain the apparent decreased catalytic efficiency in hydrolysis of the formyl group of fV bearing proteins, evidenced by their detection by PRM, and the absence of formylated methionine in this analysis (Fig. 6).

**Table 2.** ICM-Pro ligand docking scores for peptides fMAS, fVAS and fAAS.

| Peptide | Edock | Egb | Ege | Egs | Egv |
| --- | --- | --- | --- | --- | --- |
| fMAS | -32.27 | -5.96 | -12.04 | -1.57 | -61.88 |
| fVAS | -30.47 | -7.18 | -13.46 | -1.45 | -57.49 |
| fAAS | -27.73 | -7.17 | -13.08 | -0.74 | -51.01 |

**Figure 8.**
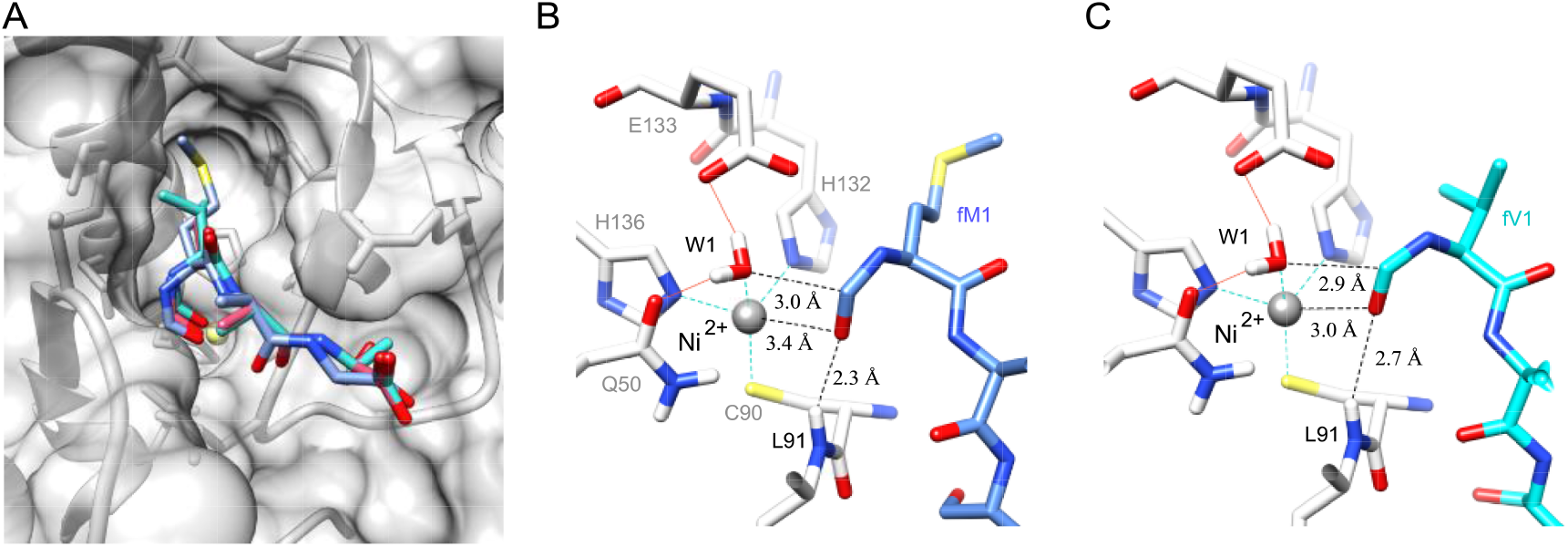
Overlay of models of highest scoring docking pose of formylated peptides fMAS, fVAS and fAAS in the substrate binding pocket of Ni^2+^ bound *E. coli* peptide deformylase. (A) A model of *E. coli* peptide deformylase (pdb code: 1bs6) is shown in grey ribbon, with the surface rendered to illustrate the peptide binding pocket (sidechains, grey sticks; active site Ni^2+^ is shown as a yellow sphere). The highest scoring docking poses for peptides fMAS (blue), fVAS (cyan) and fAAS (pink) are illustrated as sticks, with atoms coloured by type (sulphur, yellow; oxygen, red; nitrogen, blue). Figures were prepared in Chimera. (B) The active site binding interactions of the highest scoring pose of fMAS and (C) fVAS peptides, illustrating Gln50 and Glu133 hydrogen bonding to water (W1) (red lines); and nickel (Ni^2+^, grey sphere) coordination by Cys90, His132 and His136 (cyan dashed lines). Distances between modelled formyl carbon and W1, Ni^2+^ and Leu91 backbone amide are illustrated as black dashed lines.

**Figure 9.**
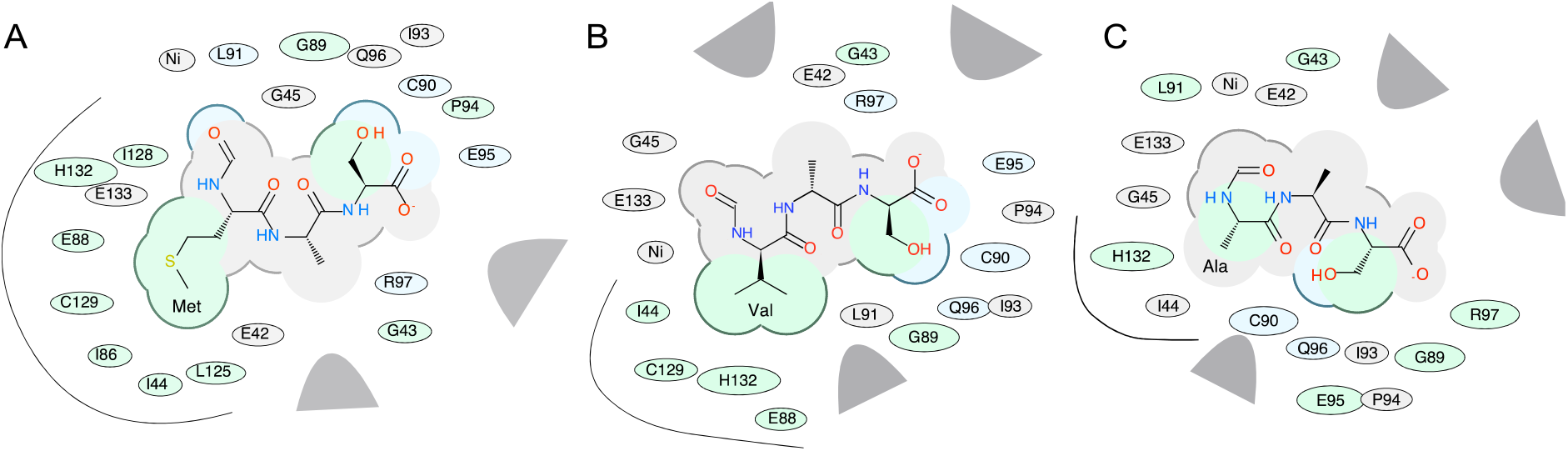
*E. coli* peptide deformylase binding pocket interactions of *in silico* docked peptides fMAS, fVAS and fAAS. 2D chemical interaction diagram for best scoring docked fMAS peptide (A), fVAS (B) and fAAS (C). The shading represents hydrophobic interactions (green), hydrogen bond acceptors (blue); van der Waals contacts (grey residues); grey parabolas represent large accessible surface areas; thick lines around ligands illustrate accessible surface; and the size of the residue oval shapes represents the strength of the contact. Black curved lines highlight the side chain interactions for residues at position 1 (Met, Val and Ala, A—C, respectively). Figures prepared in ICM-Pro.

## Discussion

In this study, we directly assessed the orthogonality of translation initiation from a mutant initiator tRNA in *E. coli* for the first time. To our knowledge, this is the first instance that the DESeq2 software was used to analyse the protein expression differences in the context of translation initiation using mutant i-tRNAs, and non-canonical start codons. Additionally, we identified the amino acids and their formylation status carried by the mutant i-tRNA(AAC). Finally, we modelled the interaction of these formylated amino acids with peptide deformylase to explain the apparent slow kinetics of deformylating valine bearing proteins.

Using high-resolution PRM mass spectrometry we clearly show there is a heterogenous population of both methionine and valine bearing N-terminal peptides on (GUU)Nluc expressed in the presence of i-tRNA(AAC) (Fig. 6). Based on past work showing that the native initiator tRNA can weakly recognize GUU start codons and initiate translation [31, 32], coupled with our detection of methionine bearing Nluc peptides in both the presence (Fig. 6C) and absence (Fig. 6B) of i-tRNA(AAC) expression, suggests this as the most likely explanation for methionine bearing Nluc peptides, not misacylation of i-tRNA(AAC) with methionine.

Despite the AAC anticodon not being present on any *E. coli* tRNAs, the valyl-tRNA synthetase (ValRS) appears to interact with i-tRNA(AAC) as if it was a valine elongator tRNA and charge it with valine. This is likely due to i-tRNA(AAC) bearing the major determinants A35 and C36 (Fig. 1C).

Additionally, i-tRNA(AAC) may also possess the minor determinant G20 motif known to enable recognition of a valyl-tRNA by ValRS [43], but complicating matters, the i-tRNA(AAC) has one additional base in its D-loop compared to the valyl-tRNA. Interestingly, it has also been shown that converting U29:A41 to C29:G41 (Fig. 1B and 1C) can act as a negative determinant and reduce aminoacylation efficiency 50-fold, likely due to an increased rigidity of the anticodon stem [43]. The presence of the 3GC motif on i-tRNA(ACC) (Fig. 1C), while permitting it to act as an initiator, increases rigidity of the anticodon stem and potentially contributes toward reducing the aminoacylation efficiency, shown by initiation rates that are approximately 100 fold lower than those from AUG start codons (Fig. 3). A further factor that may be reducing translation initiation rates from i-tRNA(AAC) is the likely requirement for valine formylation prior to productive association of the tRNA with the initiation complex [44]. *In vitro* work showed that the V_max_ of methionyl-tRNA formyltransferase formylating valine-bearing native *E. coli* initiator tRNAs was reduced 60-fold from methionine-bearing tRNAs [45]. Other work has shown that translation initiation *in vivo* with initiator tRNAs bearing valine was made more efficient by the overexpression of methionyl-tRNA formyltransferase [8], further supporting the idea that low formylation efficiency may be a significant contributor to the low initiation efficiency of i-tRNA(AAC).

Previous work that has shown proteins initiated by i-tRNAs with anticodons matching valine elongator tRNAs bear formylated N-terminal valine [8]. Using PRM mass spectrometry we readily identified formylated valine peptides from Nluc expressed in the presence of i-tRNA(AAC). N-terminal de-formylation and methionine excision occur co-translationally in *E. coli* and we wanted to understand how deformylation may be affected by a formylated-valine at the N-terminus of newly initiated proteins. Using structural modeling that compared the interaction of all possible formylated amino acids (except proline) to formylated methionine (Table 1), we found that the catalytic efficiency of hydrolysis of the formyl group attached to fVAS was likely lower in comparison to the native substrate fMAS (Figs. 8 and 9). The inability to remove the formyl group has been shown to compromise host viability [46, 47] and this would have potential implications if i-tRNA(AAC) was used more widely to control essential genes within the cell, or to express a protein that relies on quick removal of the N-terminal residue for catalytic activity [48].

Protein half-life is known to be influenced by the identity of the N-terminal amino acid and its interactions with N-degron proteins such as by ClpS discriminator proteins that transfer the bound protein to ClpAP degradation complexes [49, 50]. In theory, the inefficient removal of the formyl group from formylated-valine bearing proteins in our translation system should block recognition by ClpS and slow degradation, which was seen previously in formylated-phenylalanine initiated-proteins [8]. Surprisingly, we see the opposite effect, with i-tRNA(AAC) initiated proteins having significantly shorter *in vivo* lifetimes than methionine-bearing proteins (Fig. 7C). This feature may be useful when i-tRNA(AAC) is used as an orthogonal translation initiator if rapid protein turnover is a desirable feature, such as in biosensors [51], or whenever degradation tags might otherwise be deployed. Furthermore, the i-tRNA(AAC) initiated proteins may be used to quickly and easily relieve reliance on a single cellular degradation system, such as ClpXP, that can be overloaded [52].

## Materials and Methods

### Bacterial Strains

All plasmids within this study were expressed in BL21(DE3) (ATCC #BAA-1025) or BL21(DE3)pLysS strains (Promega, Cat. No. L1195). All strains were grown in lysogeny broth Miller (10 g/L tryptone, 10 g/L NaCl, 5 g/L yeast extract and Milli-Q® water) (LB^M^) or on LB^M^ agar (10 g/L tryptone, 10 g/L NaCl, 5 g/L yeast extract, 15 g/L agar and Milli-Q® water), supplemented with appropriate antibiotics. Chloramphenicol (25 µg/mL) was used in the media for BL21(DE3)pLysS growth, spectinomycin (50 µg/mL) for strains containing a pULTRA plasmid, and carbenicillin (100 µg/mL) for strains containing pET20b plasmids. All bacterial cultures within this study were inoculated into 2 mL of media with appropriate antibiotics in 15 mL conical falcon tubes and grown overnight (16 hours) in an Infors MT Multitron pro orbital shaker-incubator at 250 RPM rotating orbitally at 25 mm diameter at 37°C, unless stated otherwise. These stationary phase overnight liquid cultures were back diluted to 0.1 OD_600_ and grown in in 15 mL conical falcon tubes in an Infors MT Multitron pro orbital shaker-incubator at 250 RPM rotating orbitally at 25 mm diameter at 37°C, unless stated otherwise. Bacterial growth parameters have been reported to the best of our knowledge conforming with the MIEO v0.1.0 standard [53].

### Plasmid Construction

To create the pULTRA::*tac*-*metY*(AAC) plasmid we designed a modified version of *metY* from MG1655 (GenBank: U00096.3) where the anticodon region was changed from CAT to AAC (Table S4). The pULTRA-CNF plasmid (Addgene#48215) was linearized by PCR with oligos designed to flank the *metY* gene (Table S5) and the modified *metY*(AAC) gene was then inserted using homologous recombination (NEBuilder HiFi DNA Assembly kit (NEB#E2621)). The pULTRA::*tac*-Empty control plasmid was constructed through the linearization of the pULTRA::*tac*-*metY*(AAC) plasmid via PCR backbone amplification with oligos (Table S4) and ligated. This control plasmid was designed to retain all of the elements of the pULTRA::*tac*-*metY*(AAC) plasmid, except for the *metY* gene. The pET20b::T7-(NNN)*sfGFP* reporter plasmids were from previous work Hecht, Glasgow [31]. The pET20b::T7-(GUU)*Nluc6His* was built using the pET20B plasmid and *nluc* gene (Table S4). Nluc was used instead of sfGFP for mass spectrometry due to its favorable tryptic cleavage pattern.

### Bacterial Fitness Analysis

Conducted as previously Vincent, Wright [3], Vincent, Yiasemides [15] with modification. Briefly, BL21(DE3)pLysS strains containing either plasmid pULTRA::*tac*-*metY*(AAC) or pULTRA::*tac*-*Empty*, were grown overnight at 37°C with shaking in 2 mL of LB^M^ with the addition of spectinomycin and chloramphenicol. Overnight cultures of these strains were passaged and diluted 1:100 into 2 mL of fresh LB^M^ and the appropriate antibiotics and grown at 37°C with shaking until cultures were at 0.6 OD_600_. Following this, cells were back diluted to 0.1 OD_600_ with a final volume of 200 µL of LB^M^ and appropriate antibiotics in a flat bottom 96 well plate (Sigma, cat#P8116) and sealed with a gas-permeable transparent seal (Sigma cat# Z380059). Within this plate, two different conditions were set up: an induced condition containing LB^M^ with the appropriate antibiotics and 1 mM IPTG and a control condition which only contained LB^M^ and the appropriate antibiotics. Cells were grown for 7 hours at 37°C in a SPECTROStar Nano plate reader, with absorbance at OD_600_ readings occurring every 5 minutes. Growth rate (μr) and maximum cell optical density (maxOD_600_) were analyzed using the GrowthCurver R package [54].

### Fluorescence Measurements

BL21(DE3)pLysS strains containing the pULTRA::*tac*-*metY*(AAC) or the pULTRA::*tac*-Empty plasmids along with different versions of the pET20b::T7-(NNN)*sfGFP* were grown overnight at 37 °C in 2 mL of LB^M^ in the presence of carbenicillin, spectinomycin, and chloramphenicol in 96 deep well plates (Sigma, cat#P8116). Bacterial cultures were then diluted 1:100 in 400 µL of fresh LB^M^ and appropriate antibiotics and grown for 1 hour at 37 °C with shaking. After one hour of growth, the cultures were induced with 1 mM IPTG, and growth for a further 4 hours at 37 °C, or until cells reached 0.6 OD_600_, in preparation for bulk fluorescence measurements and flow cytometry.

Following induction, 400 µL of culture was transferred into a clear 96-well plate (Sigma, cat#CLS3610) and centrifuged in a swinging bucket rotor at 2,240 x*g* for 12 minutes at 20°C. Following this the supernatant was removed, and the pelleted cells were resuspended in 200 µL of phosphate-buffered saline (PBS) and transferred into a black clear bottom 96-well plates (Sigma, cat#CL3603). Resuspended cells were stored at 4°C overnight to allow for sfGFP maturation [31]. Absorbance at OD_600_ and bulk fluorescence for sfGFP expression was measured using the PHERAstar microplate reader (BMG Labtech) (excitation 488 nm, emission 530 nm). Results were recorded with a gain setting of 200. Normalized (Arb. fluorescence units/OD_600_) results were analyzed using the DEseq2 R package. Parameters were set to find fluorescence changes of 4-fold or more, with a statistical significance of p-value <0.01.

### Flow Cytometry

Bacterial cultures were prepared as for fluorescence measurements above. Following this, cells were diluted 100-fold in PBS to reach the final cell concentration of approximately 10^6^ cfu/mL and measured on a CytoFLEX S (Beckman Coulter) using a FITC fluorescence channel with a 488 nm excitation laser and 525/40 nm emission band-pass filter with side scatter triggered events threshold set to 10,000 events. Data processing and analysis was performed using CytExpert (Version 2.3.0.84) (Beckman Coulter) and the DEseq2 R package [34]. Parameters were set to find fluorescence changes of 4-fold or more, with a statical significance of *p<0*.*01*.

### Expression and Purification of Nluc

Biological triplicates of single colony BL21(DE3)pLysS cells harboring the pULTRA::*tac*-Empty or pULTRA::*tac*-*metY*(AAC) plasmids and the pET20b::T7-(AUG)*Nluc*6XHis or pET20b::T7-(GUU)*Nluc*6XHis plasmids were used to inoculate 5 mL of LB^M^ supplemented with spectinomycin, chloramphenicol, and carbenicillin and incubated at 37°C overnight with shaking. The protocol used for expression and purification of Nluc Hecht, Glasgow [31] was adopted with the following modifications. The overnight cultures were diluted 1:100 in 30 mL of LB^M^ with appropriate antibiotics and incubated at 37°C for 1 hour, followed by induction with 1 mM of IPTG and a final outgrowth step for an additional 6 hours to allow for sufficient protein expression and production.

The cell cultures were pelleted at 3,500 x*g* for 10 min. The supernatant was discarded, and the cell pellets were chemically lysed using 0.4 mL lysis buffer (CelLytic B cell lysis reagent (Sigma-Aldrich #B7310), 0.1 mg lysozyme, 1x protease inhibitor (Roche #04693132001), and 100 units of benzonase (Sigma-Aldrich #E1014)). The resuspended cells were incubated at room temperature for 20 min to allow for sufficient lysis. The soluble protein fraction was then collected by centrifuging the cell lysate at 16,000 x*g* for 10 minutes. The expressed C-terminal 6xHis-tagged Nluc was Ni-NTA purified using HisPurTMNi-NTA Resin (ThermoFisher scientific, #88221) as per the HisPurTMNi-NTA Resin protocol. Equilibration, wash, and elution buffers were made using 300 mM PBS with 20 mM, 60 mM, and 500 mM imidazole, respectively.

### Mass Spectrometry

Purified Nluc samples were reduced through the addition of dithiothreitol (DTT) to a final concentration of 10 mM and incubated on a heat block at 60°C for 30 minutes. Samples were alkylated using iodoacetamide at a final concentration of 30 mM and incubated at room temperature in the dark for 1 hour. Purified Nluc was precipitated out of solution using 1 mL of ice-cold acetone incubated overnight at -20°C and pelleted via centrifugation at 16,000 x*g* for 10 min at 4°C. The resulting protein pellet was resuspended in 8 M Urea, 50 mM Tris-HCl buffer (pH 8.0) and diluted 5-fold by addition of 50 mM Tris-HCl buffer (pH 8.0). Protein samples were digested with 1:100 (w/w) trypsin/protein incubated at 37°C overnight, then digested again for 4 hours with fresh trypsin at 37°C (in the same amounts as previously). Peptides were then acidified with 1% (v/v) formic acid, followed by a C18 stage tip clean up and filtration through glass fiber filter circles (Fisher Scientific, #09-804-24A). The peptides were then dried in a SpeedVac vacuum concentrator and resuspended to a final concentration of 1 μg/μL in 2% (v/v) acetonitrile / 0.1% (v/v) formic acid and used for PRM analysis [55]. Tryptic peptides were analyzed using an Easy-nLC1000 liquid chromatography system coupled to a high-resolution Q-Exactive mass spectrometer (Thermo Fisher Scientific).

The PRM analysis method was adopted from previous work Vincent, Wright [3].

Reconstituted peptides were injected onto a C18 reverse phase column and eluted over a 120-minute linear gradient by increasing concentrations of elution buffer (99.9% (v/v) acetonitrile, 0.1% formic acid). An initial survey scan was conducted at 70,000 resolution with a scan range of 320 to 1,800 *m/z*, AGC target set to 3 × 10^6^, and a maximum injection time of 100 ms. An unscheduled inclusion list for N-terminal peptides (File S1) was generated using Skyline [56](Version 20.1.0.155). Predefined target precursors that were identified were subjected to high collision dissociation fragmentation using a normalized collision energy of 27. Resulting product ions were analyzed in an MS/MS scan set at 17,500 resolution, AGC target of 3 × 10^6^, maximum injection time of 60 ms, and a 2.0 m/z isolation window. Analysis of precursor and product ion spectra were analyzed using MaxQuant (Version 1.6.17.0)[57] and Skyline (Version 20.1.0.155)[56].

### Protein Stability Assay

BL21(DE3)pLysS strains harboring pULTRA and pET20b Nluc reporter plasmids were picked from single colonies in biological triplicates and inoculated overnight with shaking in 5 mL of LB^M^ supplemented with appropriate antibiotics. Overnight cultures were diluted 1:100 into 15 mL of fresh LB^M^ and grown at 37°C with shaking until all cultures reached mid-log phase (0.6 OD_600_). The cultures were then subsequently back diluted to 0.1 OD_600_ into 15 mL fresh LB ^M^ and in 50 mL Falcon tubes. The cells were grown for 2 hours at 37°C with shaking prior to induction with 1 mM IPTG. After 1 hour of induction, the cells were pelleted by centrifugation at 2,500 x*g* for 10 min. In order to remove all of the IPTG in the media, the cells were washed with fresh LB^M^, and re-pelleted.

Following this, the cells were resuspended in 15 mL fresh LB^M^, and the addition of 2% glucose, to repress the production of both tRNA and Nluc. The cells were then grown for 12 hours at 37°C with shaking. Luminescence was measured from 100 μL of culture from each time point, that was mixed with 100 μL of pre-prepared Nano-Glo® Luciferase Assay Buffer + Substrate (Promega # N1130) and incubated for 20 minutes at room temperature. Following incubation, luminescence was measured using a Synergy H1 Hybrid Multi-Mode Reader at 100 gain setting. Arbitrary luminescence units were then normalized by the absorbance (OD_600_) measurement of the culture at the point of sampling.

### Protein:protein Interaction Stability and Docking Analyses

Peptide binding stability and optimal docking of substrate tripeptides were calculated in Molsoft ICM version 3.9-2a [58] using the pdb-redo [59] refined structure of *E. coli* peptide deformylase 1bs6. The structure of peptide deformylase was cleaned of unnecessary small molecules and converted to an ICM object using an object conversion protocol that incorporates addition of hydrogens, optimisation of His, Asn and Gln rotamers, and assignment of protonation states and secondary structure [58].

The model of chains A (enzyme) and D (peptide MAS) was prepared by global optimisation of side chains and backbone annealing in ICM-Pro. Waters W1 and W2 (as defined by [36]) (waters 2041 and 2042, 1bs6), and Ni^2+^ were maintained in the active site for peptide binding stability calculations. Met1 was mutated to all side chains (except Pro) and relative protein:protein binding free energy ΔΔG calculated for all mutants [38]. Formylated peptides fMAS, fVAS and fAAS were built using the ICM-Pro ligand editor and energy minimised [60]. Formylated peptides were docked into the enzyme active site of 1bs6 chain A including W1 and Ni^2+^ using flexible ligand docking with relaxed covalent geometry, automatic assignment of charged groups, and fixed peptide-like amide bonds. The binding site was defined based on the position of the bound MAS peptide (1bs6, chain D) in the original structure (1bs6, chain A). Docking scores were calculated in ICM-Pro with docking effort of 5 and the top ten conformers stored for analysis of optimal binding poses [58]. 2D interaction diagrams were prepared in ICM-Pro and structural model figures were prepared with Chimera [61].

## Acknowledgements

We recognize that this research was conducted on the traditional lands of the Wattamattagal clan of the Darug nation. We thank Ariel Hecht for providing R code for the production of several figures, Daniel McDougal for helpful discussions and feedback on the manuscript, and the Australian Proteome Analysis Facility. AH was supported by a Macquarie Research Excellence PhD Scholarship and CSIRO SynBio FSP Top-Up scholarship, DS supported by a Macquarie Research Excellence PhD Scholarship, FW was supported by the Ramsay Fellowship of Applied Science, and PRJ was supported by the Molecular Sciences Department, Faculty of Science & Engineering, and the Deputy Vice-Chancellor (Research) of Macquarie University.

## Author Contributions

**Andras Hutvagner**: Conceptualization, Experimentation, Formal Analysis, Investigation, Methodology, Visualization, Writing - Original Draft, Writing - Review, Funding acquisition. **Dominic Scopelliti**: Conceptualization, Experimentation, Formal Analysis, Investigation, Methodology, Visualization, Writing - Original Draft, Writing - Review.

**Fiona Whelan:** Conceptualization, Formal analysis, Investigation, Methodology, Visualization, Writing - Original Draft, Writing - Review.

**Paul R. Jaschke**: Conceptualization, Writing - Review & Editing, Supervision, Project administration, Funding acquisition.

