## Supplementary material for "Orthogonal translation initiation using the non-canonical initiator tRNA(AAC) alters protein sequence and stability *in vivo*": Suplementary Information

**CONTENTS**

**Supporting Figures**

Figure S1	Flow cytometry data of pULTRA:*tac:metY*(AAC) and pULTRA:Empty strains expressing sfGFP from all 64 start codons.

Figure S2	Parallel reaction monitoring detection of Nluc reporter proteins bearing a N-	terminal methionine from BL21(DE3)pLysS (pULTRA::*tac*-*metY*(AAC);pET20b:T7-(GUU)*Nluc*6HIS).

Figure S3	Parallel reaction monitoring detection of Nluc reporter proteins bearing a N-	terminal formylated methionine from BL21(DE3)pLysS (pULTRA::*tac*-*Empty*;pET20b:T7-(AUG)*Nluc*6HIS).

**Supporting Tables**

Table S1	ICM-Pro peptide docking scores fMAS

Table S2	ICM-Pro peptide docking scores fVAS

Table S3	ICM-Pro peptide docking scores fAAS

Table S4	Gene Sequences

Table S5	Oligos used (sequencing, Gibson assembly, colony PCR for all plasmids)

**Supporting Files**

Supporting File S1:	PRM inclusion list

Supporting File S2: 	Bulk fluorescence data

Supporting File S3: 	Differential expression analysis

Supporting File S4: 	Fitness Assay

Supporting File S5:	Plasmid sequence: pULTRA:*tac:metY*(AAC)

Supporting File S6:	Plasmid sequence: pet20b:*t7:sfGFP*(NNN)

Supporting File S7:	Plasmid sequence pet20b:*t7:nluc*(GTT)

Supporting File S8:	Plasmid sequence pet20b:*t7:nluc*(ATG)

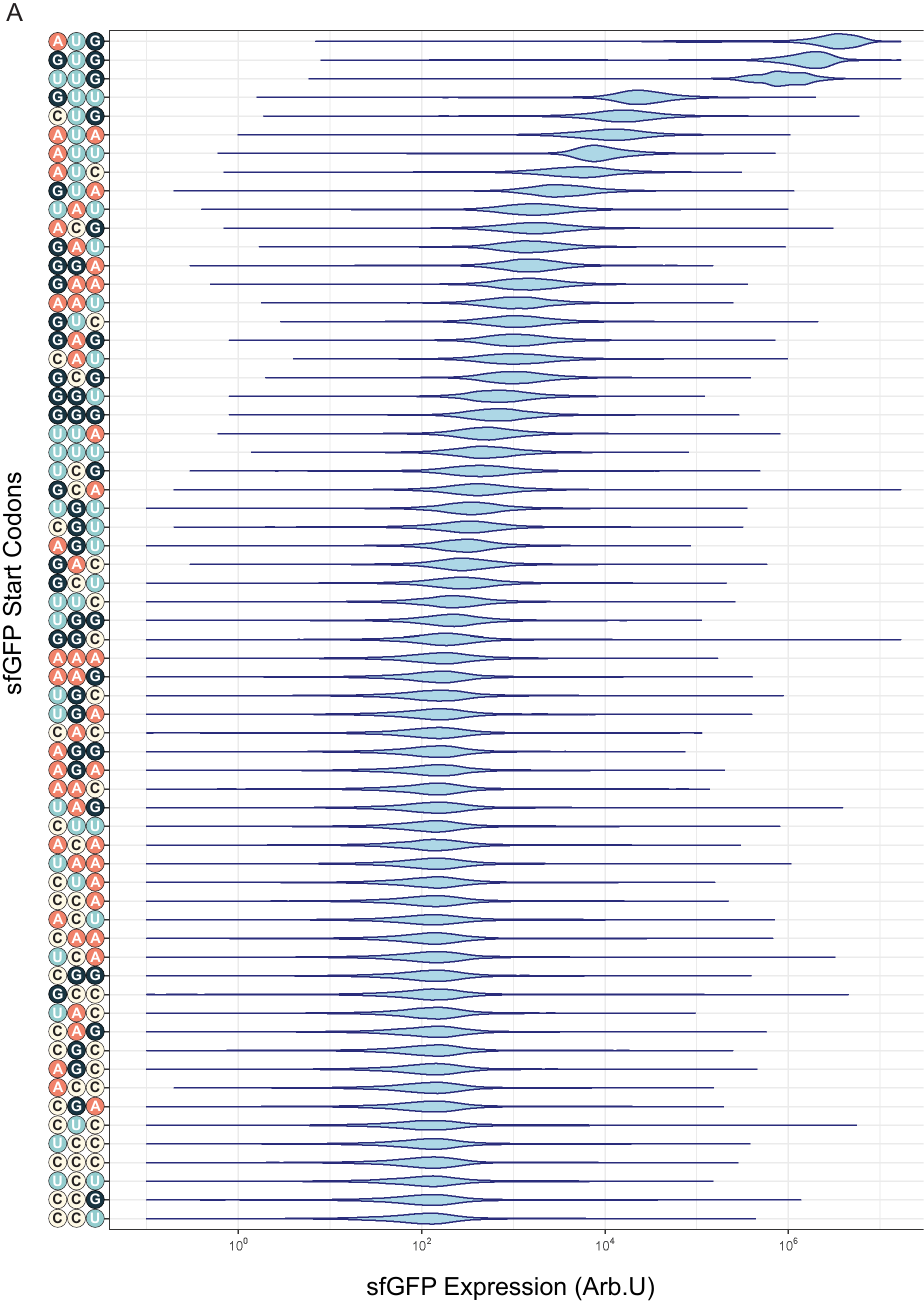

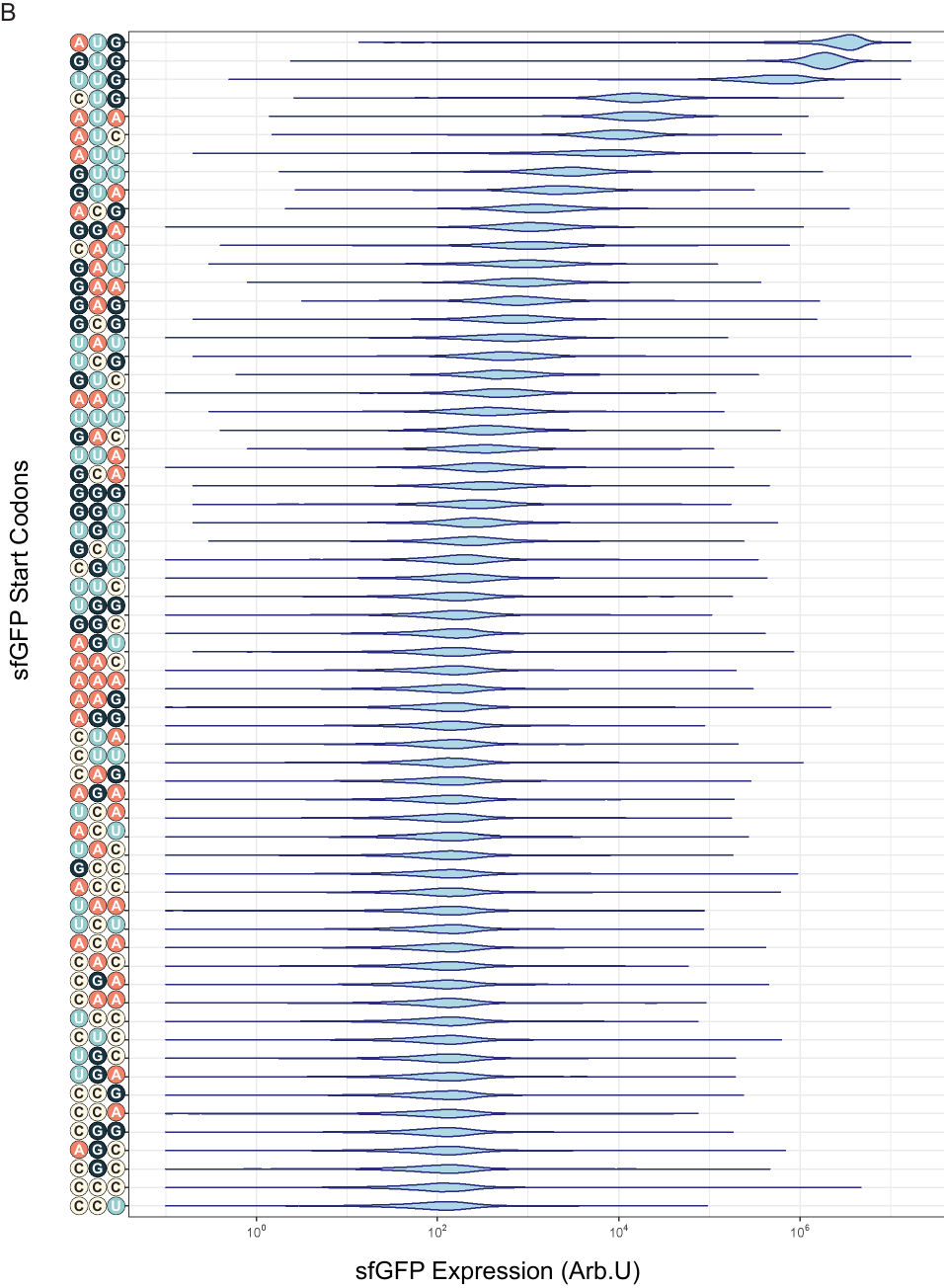
**Fig. S1 Violin Plots of Flow Cytometry Data.** Violin plots of the flow cytometry data for both (A) pULTRA:*tac*:*metY*(AAC) and (B) pULTRA:*tac*:Empty strains, expressing sfGFP from all possible 64 start codons show unimodal distribution. Cells were measured on a CytoFLEX S (Beckman Coulter) using a FITC fluorescence channel with a 488 nm excitation laser and 525/40 nm emission band-pass filter with side scatter triggered events threshold set to 10,000 events.

­­­
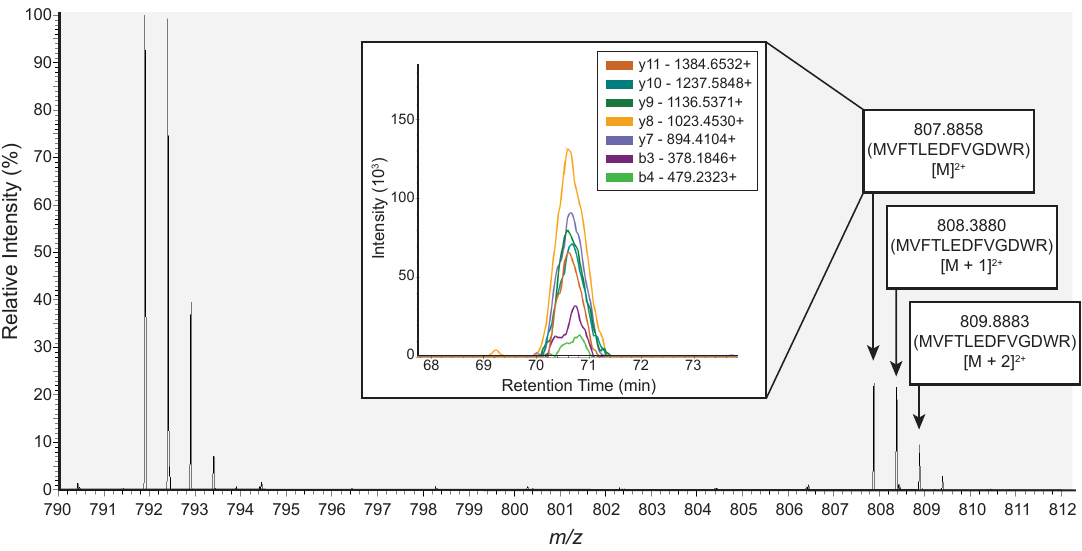

**Fig. S2 PRM detection of methionine bearing N-terminal peptides purified from *E. coli* BL21(DE3)pLysS (pULTRA::*tac*-*metY*(AAC); pET20b:*T7*-(GUU)*Nluc*6HIS).** MS1 scan during retention time range of 70-73 minutes of Nluc reporter protein expressed in BL21(DE3)pLysS (pULTRA::*tac*-*metY*(AAC); pET20b:*T7*-(GUU)*Nluc*6HIS). Labelled precursor ion peaks (807.8858 *m/z*) consistent with an N-terminal peptide bearing methionine (MVFTLEDFVGDWR) and their respective isotopes at the N-terminus are shown. The inset (window coming off of MS1 signal label), shows MS2 y^-^ and b^-^ series product ion signals used to confirm amino acid sequence of N-terminal peptide. Nluc6His reporter proteins expressed in BL21(DE3)pLysS were induced with 1mM IPTG.­­­

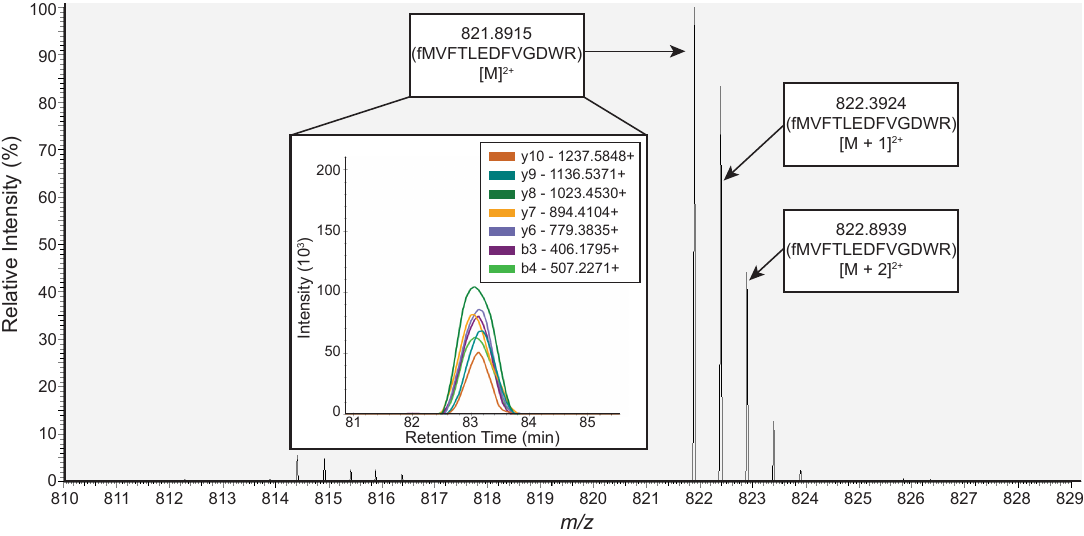

**Fig. S3 PRM detection of formylated methionine bearing N-terminal peptides purified from *E. coli* BL21(DE3)pLysS (pULTRA::*tac*-Empty; pET20b:*T7*-(AUG)*Nluc*6HIS).** MS1 scan during retention time range of 81-85 minutes of Nluc reporter protein expressed in BL21(DE3)pLysS (pULTRA::*tac*-*metY*(AAC); pET20b:*T7*-(AUG)*Nluc*6HIS). Labelled precursor ion peaks (821.8915 *m/z*) consistent with an N-terminal peptide bearing formylated methionine (fMVFTLEDFVGDWR) and their respective isotopes at the N-terminus are shown. The inset (window coming off of MS1 signal label), shows MS2 y^-^ and b^-^ series product ion signals used to confirm amino acid sequence of N-terminal peptide. Nluc6His reporter proteins expressed in BL21(DE3)pLysS were induced with 1mM IPTG.­­­

**SUPPORTING TABLES**

**Table S1 ICM-Pro peptide docking scores fMAS**

| *Peptide* | Edock | Egb | Ege | Egs | Egv |
| --- | --- | --- | --- | --- | --- |
| *fMAS_1* | -32.27 | -5.96 | -12.04 | -1.57 | -61.88 |
| *fMAS_2* | -29.59 | -11.80 | -16.35 | -1.23 | -53.45 |
| *fMAS_3* | -25.87 | -6.70 | -10.40 | -1.56 | -58.50 |
| *fMAS_4* | -24.30 | -10.39 | -11.58 | -1.40 | -53.61 |
| *fMAS_5* | -24.10 | -8.28 | -12.37 | -2.29 | -52.42 |
| *fMAS_6* | -22.37 | -7.99 | -5.18 | -0.58 | -62.31 |
| *fMAS_7* | -21.70 | -8.04 | -7.70 | -0.54 | -59.63 |
| *fMAS_8* | -21.27 | -14.27 | -13.02 | -1.53 | -43.71 |
| *fMAS_9* | -21.23 | -7.02 | -12.91 | -1.81 | -52.36 |
| *fMAS_10* | -21.19 | -14.31 | -14.61 | -1.35 | -43.70 |

**Table S2 ICM-Pro peptide docking scores fVAS**

| *Peptide* | Edock | Egb | Ege | Egs | Egv |
| --- | --- | --- | --- | --- | --- |
| *fVAS_1* | -30.47 | -7.18 | -13.46 | -1.45 | -57.49 |
| *fVAS_2* | -28.39 | -9.31 | -7.45 | -0.22 | -57.17 |
| *fVAS_3* | -26.58 | -10.57 | -12.31 | -1.53 | -48.77 |
| *fVAS_4* | -25.85 | -12.77 | -16.82 | -1.42 | -46.69 |
| *fVAS_5* | -22.88 | -13.63 | -16.08 | -1.82 | -40.81 |
| *fVAS_6* | -22.82 | -13.18 | -17.79 | -1.56 | -38.50 |
| *fVAS_7* | -20.78 | -13.57 | -17.80 | -1.41 | -31.60 |
| *fVAS_8* | -20.39 | -7.85 | -9.13 | 1.03 | -52.82 |
| *fVAS_9* | -20.20 | -10.46 | -5.61 | -0.39 | -51.24 |
| *fVAS_10* | -20.14 | -7.59 | -3.89 | 0.42 | -58.84 |

**Table S3 ICM-Pro peptide docking scores fAAS**

| *Peptide* | Edock | Egb | Ege | Egs | Egv |
| --- | --- | --- | --- | --- | --- |
| *fAAS_1* | -27.73 | -7.17 | -13.08 | -0.74 | -51.01 |
| *fAAS_2* | -26.59 | -14.01 | -16.82 | -1.35 | -37.20 |
| *fAAS_3* | -24.47 | -11.14 | -14.71 | -0.76 | -42.66 |
| *fAAS_4* | -21.45 | -13.48 | -18.37 | -0.82 | -30.11 |
| *fAAS_5* | -20.45 | -11.08 | -7.38 | -0.56 | -49.25 |
| *fAAS_6* | -19.87 | -7.49 | -10.14 | -0.94 | -46.93 |
| *fAAS_7* | -19.83 | -6.96 | -8.48 | -1.66 | -47.87 |
| *fAAS_8* | -19.42 | -9.27 | -5.51 | -0.41 | -49.30 |
| *fAAS_9* | -19.15 | -8.64 | -12.07 | -0.85 | -40.53 |
| *fAAS_10* | -19.07 | -7.50 | -5.60 | -0.15 | -53.65 |

**Table S4: Gene sequences**

| Gene | Sequence (5’-3’) |
| --- | --- |
| *sfgfp* | ATGCGTAAAGGCGAAGAGCTGTTCACTGGTGTCGTCCCTATTCTGGTGGAACTGGATGGTGATGTCAACGGTCATAAGTTTTCCGTGCGTGGCGAGGGTGAAGGTGACGCAACTAATGGTAAACTGACGCTGAAGTTCATCTGTACTACTGGTAAACTGCCGGTACCTTGGCCGACTCTGGTAACGACGCTGACTTATGGTGTTCAGTGCTTTGCTCGTTATCCGGACCATATGAAGCAGCATGACTTCTTCAAGTCCGCCATGCCGGAAGGCTATGTGCAGGAACGCACGATTTCCTTTAAGGATGACGGCACGTACAAAACGCGTGCGGAAGTGAAATTTGAAGGCGATACCCTGGTAAACCGCATTGAGCTGAAAGGCATTGACTTTAAAGAAGACGGCAATATCCTGGGCCATAAGCTGGAATACAATTTTAACAGCCACAATGTTTACATCACCGCCGATAAACAAAAAAATGGCATTAAAGCGAATTTTAAAATTCGCCACAACGTGGAGGATGGCAGCGTGCAGCTGGCTGATCACTACCAGCAAAACACTCCAATCGGTGATGGTCCTGTTCTGCTGCCAGACAATCACTATCTGAGCACGCAAAGCGTTCTGTCTAAAGATCCGAACGAGAAACGCGATCATATGGTTCTGCTGGAGTTCGTAACCGCAGCGGGCATCACGCATGGTATGGATGAACTGTACAAATGATGA |
| *Nluc(6-HIS)* | ATGGTCTTCACACTCGAAGATTTCGTTGGGGACTGGCGACAGACAGCCGGCTACAACCTGGACCAAGTCCTTGAACAGGGAGGTGTGTCCAGTTTGTTTCAGAATCTCGGGGTGTCCGTAACTCCGATCCAAAGGATTGTCCTGAGCGGTGAAAATGGGCTGAAGATCGACATCCATGTCATCATCCCGTATGAAGGTCTGAGCGGCGACCAAATGGGCCAGATCGAAAAAATTTTTAAGGTGGTGTACCCTGTGGATGATCATCACTTTAAGGTGATCCTGCACTATGGCACACTGGTAATCGACGGGGTTACGCCGAACATGATCGACTATTTCGGACGGCCGTATGAAGGCATCGCCGTGTTCGACGGCAAAAAGATCACTGTAACAGGGACCCTGTGGAACGGCAACAAAATTATCGACGAGCGCCTGATCAACCCCGACGGCTCCCTGCTGTTCCGAGTAACCATCAACGGAGTGACCGGCTGGCGGCTGTGCGAACGCATTCTGGCGCACCACCACCACCACCAC |
| *metY(AAC)* | CGCGGGGTGGAGCAGCCTGGTAGCTCGTCGGGCTAACAACCCGAAGATCGTCGGTTCAAATCCGGCCCCCGCAACCA |

**Table. S5: Oligos used (sequencing, Gibson assembly, colony PCR for all plasmids)**

| Oligo | Sequence (5’-3’) |
| --- | --- |
| *metY* Sequencing and colony PCR Primer Forward | TCTCCCTTATGCGACTCCTG |
| *metY* Sequencing and colony PCR primer Reverse | AGATCCGGCCACGATGAC |
| *sfgfp* Sequencing and colony PCR Primer Forward | CTCGCGTATCGGTGATTCAT |
| *sfgfp* Sequencing and colony PCR Primer Reverse | CGGATAACGAGCAAAGCACT |
| *pULTRA* linearisation primer Forward | CGCAACCACTTTCCCTTAGA |
| *pULTRA* linearisation primer Reverse | TGAAAGCACCTCCTTTGTGA |
| *nluc* Sequencing and colony PCR Primer Forward | CGCGTTTCCAGACTTTACG |
| *nluc* Sequencing and colony PCR Primer Reverse | GCTTAATGCGCCGCTACAG |
